# Evolutionarily-conserved chromatin crosstalk generates a DOT1L-dose dependency in thymic lymphoma caused by loss of HDAC1

**DOI:** 10.1101/509976

**Authors:** Hanneke Vlaming, Chelsea M. McLean, Tessy Korthout, Mir Farshid Alemdehy, Sjoerd Hendriks, Cesare Lancini, Sander Palit, Sjoerd Klarenbeek, Eliza Mari Kwesi-Maliepaard, Thom M. Molenaar, Liesbeth Hoekman, Thierry T. Schmidlin, A.F. Maarten Altelaar, Tibor van Welsem, Jan-Hermen Dannenberg, Heinz Jacobs, Fred van Leeuwen

**Affiliations:** Division of Gene Regulation, Netherlands Cancer Institute, Amsterdam, The Netherlands; Division of Tumor Biology & Immunology, Netherlands Cancer Institute, Amsterdam, The Netherlands; Experimental Animal Pathology, Netherlands Cancer Institute, Amsterdam, The Netherlands; Proteomics Facility, Netherlands Cancer Institute, Amsterdam, The Netherlands; Biomolecular Mass Spectrometry and Proteomics, Bijvoet Center for Biomolecular Research and Utrecht Institute for Pharmaceutical Sciences, Utrecht University and Netherlands Proteomics Centre, Utrecht, The Netherlands; Current address: Department of Biological Chemistry and Molecular Pharmacology, Harvard Medical School, Boston, MA, USA; Current address: Genmab B.V., Antibody Sciences, Utrecht, The Netherlands.

**Author notes:** These authors contributed equally to this work (Hanneke Vlaming, Chelsea M. McLean).

**Keywords:** DOT1L, Dot1, HDAC1, H3K79, histone methylation, histone acetylation, histone ubiquitination, lymphoma

## Abstract

DOT1L methylates histone H3K79 and is aberrantly regulated in MLL-rearranged leukemia. Inhibitors have been developed to target DOT1L activity in leukemia but the cellular mechanisms that regulate DOT1L are still poorly understood. Here we identify the budding yeast histone deacetylase Rpd3 as a negative regulator of Dot1. At its target genes, the transcriptional repressor Rpd3 restricts H3K79 methylation, explaining the absence of H3K79me3 at a subset of genes in the yeast genome. Similar to the crosstalk in yeast, inactivation of the murine Rpd3 homolog HDAC1 in thymocytes led to an increase in H3K79 methylation. Thymic lymphomas that arise upon genetic deletion of *Hdac1* retained the increased H3K79 methylation and were sensitive to reduced DOT1L dosage. Furthermore, cell lines derived from *Hdac1^Δ/Δ^* thymic lymphomas were sensitive to DOT1L inhibitor, which induced apoptosis. In summary, we identified an evolutionarily-conserved crosstalk between HDAC1 and DOT1L with impact in murine thymic lymphoma development.

## Introduction

Aberrant histone modification patterns have been observed in many diseases and this deregulation of chromatin can play a causative role in disease. Since epigenetic alterations are, in principle, reversible in nature, histone (de)modifiers are attractive therapeutic targets (Brien et al. 2016; Jones et al. 2016; Shortt et al. 2017). Several epigenetic drugs are currently in the clinic or in clinical trials, but for many of the drug targets we are only beginning to understand their cellular regulation.

The histone H3K79 methyltransferase DOT1L (KMT4; Dot1 in yeast) is an epigenetic enzyme for which inhibitors are in clinical development for the treatment of MLL-rearranged (MLL-r) leukemia (Stein and Tallman 2016). In MLL-r leukemia, DOT1L recruitment to MLL target genes, such as the HoxA cluster leads to aberrant H3K79 methylation and increased transcription (reviewed in Vlaming and Van Leeuwen 2016). Although the DOT1L inhibitor Pinometostat (EPZ-5676) has shown promising results in the lab and is currently in clinical development (Bernt et al. 2011; Daigle et al. 2013; Waters et al. 2015; Stein and Tallman 2016; Stein et al. 2018), the cellular mechanisms and consequences of DOT1L deregulation are only just being uncovered (Vlaming and Van Leeuwen 2016).

An important mechanism of regulation is the trans-histone crosstalk between monoubiquitination of the C terminus of histone H2B (H2Bub) at lysine 120 (123 in yeast) and methylation of histone H3K79 (Zhang et al. 2015). The addition of a ubiquitin peptide to the nucleosome at this position occurs in a co-transcriptional manner and promotes the activity of Dot1/DOT1L, possibly by activation of DOT1L or coaching it towards H3K79 and thereby increasing the chance of a productive encounter (Zhou et al. 2016; Vlaming et al. 2014). Another mechanism of regulation is mediated by the direct interactions of DOT1L with central transcription elongation proteins (reviewed in Vlaming and Van Leeuwen 2016). These interactions target DOT1L to transcribed chromatin and provide an explanation for the aberrant recruitment of DOT1L by oncogenic MLL fusion proteins (Deshpande et al. 2014; Chen et al. 2015; Li et al. 2014; Kuntimaddi et al. 2015; Wood et al. 2018). Further characterizing the regulatory network of DOT1L could lead to the identification of alternative drug targets for diseases in which DOT1L is critical and provide alternative strategies in case of resistance to treatment with DOT1L inhibitors (Campbell et al. 2017).

In a previous study, we presented a ChIP-barcode-seq screen (Epi-ID) identifying novel regulators of H3K79 methylation in yeast (Vlaming et al. 2016). The Rpd3-large (Rpd3L) complex was identified as an enriched complex among the candidate negative regulators of H3K79 methylation of a barcoded reporter gene. Rpd3 is a class I histone deacetylase (HDAC) that removes acetyl groups of histones, as well as numerous non-histone proteins, and is generally associated with transcriptional repression (Yang and Seto 2008). Several inhibitors of mammalian HDACs have been approved for the treatment of cutaneous T-cell lymphoma and other hematologic malignancies, while others are currently being tested in clinical trials (West and Johnstone 2014). HDAC1 and HDAC2, prominent members of the class I HDACs, are found in the repressive Sin3, NuRD, and CoREST complexes (Yang and Seto 2008). Loss or inhibition of HDAC1/Rpd3 leads to increased histone acetylation, which in turn can lead to increased expression of target genes and cryptic transcripts (Carrozza et al. 2005; Joshi and Struhl 2005; Li et al. 2007; Rando and Winston 2012; McDaniel and Strahl 2017; Brocks et al. 2017).

Here, we demonstrate that Rpd3 restricts H3K79 methylation at its target genes. Most euchromatic genes in the yeast genome are marked by high levels of H3K79me3. We observed that a subset of the genes that do not follow this pattern has lower H3K79me3 levels due to the action of the Rpd3L complex, which deacetylates its targets and imposes strong transcriptional repression and absence of H2Bub1. Importantly, the Rpd3-Dot1 crosstalk is conserved in mammals: genetic ablation of *Hdac1* in murine thymocytes also leads to an increase in H3K79 methylation *in vivo*. High H3K79me is maintained in the lymphomas these mice develop, and a reduction in DOT1L activity by heterozygous deletion of *Dot1L* reduces tumor burden, an effect that was not observed upon homozygous deletion of *Dot1L*. Furthermore, DOT1L inhibitors induce apoptosis in *Hdac1*-deficient but not *Hdac1*-proficient thymic lymphoma cell lines, suggesting a DOT1L-dose dependence. Taken together, our studies reveal a new, evolutionarily-conserved mechanism of H3K79me regulation by Rpd3/HDAC1 with relevance for cancer development.

## Results

### Identification of the Rpd3L complex as a negative regulator of H3K79 methylation

We recently reported a systematic screening strategy called Epi-ID to identify regulators of H3K79 methylation (Vlaming et al. 2016). In that screen, relative H3K79 methylation (H3K79me) levels at two DNA barcodes (UpTag and DownTag) flanking a reporter gene were measured in a genome-wide library of barcoded deletion mutants, thus testing thousands of genes for H3K79me regulator activity at these loci (Fig. 1A). Since higher Dot1 activity in yeast leads to a shift from lower (me1) to higher (me3) methylation states (Frederiks et al. 2008), the H3K79me3 over H3K79me1 ratio was used as a measure for Dot1 activity. A growth-corrected H3K79me score was calculated to account for the effect of growth on H3K79 methylation and groups of positive and negative candidate regulators were identified (Vlaming et al. 2016). Components of the Rpd3L complex were enriched among candidate negative regulators (10-fold over-representation, p=1.2E-4) (Vlaming et al. 2016). The histone deacetylase Rpd3 is found in two complexes, the large Rpd3L complex and the small Rpd3S complex, which also share the subunits Sin3 and Ume1 (Carrozza et al. 2005; Keogh et al. 2005). A closer inspection of the Rpd3 complexes revealed that deletion of Rpd3L subunits resulted in an increase in H3K79 methylation on both the UpTag and DownTag (promoter and terminator context, respectively; Fig. 1B), with the exception of two accessory subunits that play peripheral roles (Lenstra et al. 2011). Deletion of the two Rpd3S-specific subunits did not lead to an increase in H3K79me (Fig. 1B), which is consistent with Rpd3S binding and acting on coding sequences (Drouin et al. 2010) and thus away from the intergenic barcodes. To validate the effect on a global scale, we performed targeted mass spectrometry analysis to determine the relative levels of the different H3K79me states (me0 to me3) in *rpd3Δ* and *sin3Δ* strains. On bulk histones, these strains showed an increase in H3K79me (increase in H3K79me3 at the cost of lower methylation states; Fig. 1C). The H3K79me increase was not caused by an increase in Dot1 protein (Supplemental Fig. 1A) or mRNA expression (Kemmeren et al. 2014). Thus, although these regulators were identified using only two 20-base-pair barcodes to read out H3K79me levels at a reporter locus, their effects could be validated globally.

**Figure 1.**
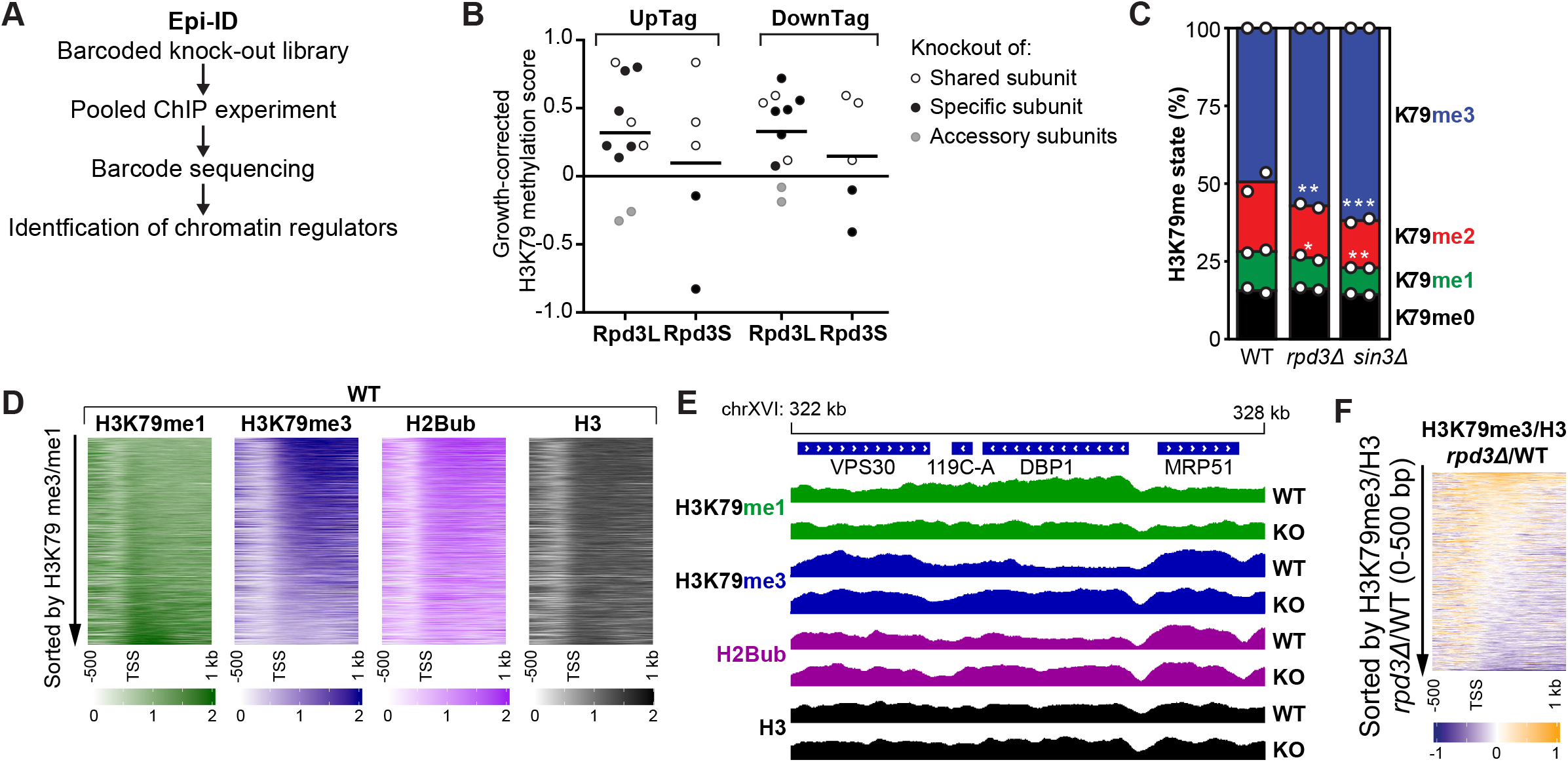
Rpd3 and other members of the Rpd3L complex negatively regulate H3K79 methylation. A) Schematic overview of the Epi-ID strategy. B) Epi-ID H3K79 methylation scores of the deletion mutants of members of the Rpd3L and Rpd3S complexes, calculated as described in the Supplemental Methods, where 0 means a wild-type H3K79me level (log2 scale). The gray dots indicate accessory subunits. UpTag and DownTag are barcode reporters in a promoter and terminator context, respectively. Data was obtained on all Rpd3L/Rpd3S subunits except Sds3. C) Mass spectrometry analysis of H3K79 methylation in wild-type and mutant strains. Mean and individual data points of two biological replicates. D) Heatmaps of H3K79me1, H3K79me3, H2Bub and H3 in wild-type cells, aligned on the TSS. Genes were sorted based on the average H3K79me3/H3K79me1 ratio in the first 500 bp. E) Snapshot of depth-normalized ChIP-seq data tracks from wild-type and *rpd3Δ* strains showing 6 kb surrounding the *DBP1* ORF, which is the top gene in the heatmap in panel F. All tracks have the same y axis (0-2). A snapshot of another top-regulated gene is shown in Supplemental Fig. 1D. F) Heatmap of the H3K79me3/H3 change in *rpd3Δ* versus wild-type cells, aligned on the TSS. Genes were sorted based on the average ratio in the first 500 bp.

### Rpd3 represses H3K79 methylation at the 5’ ends of a subset of genes

We next asked at which regions Rpd3 and Sin3 regulate H3K79 methylation in yeast, other than the barcoded reporter gene. To address this, we performed ChIP-seq analysis for H3K79me1, H3K79me3, and H3 in wild-type and *rpd3Δ* strains. In addition, we included ChIP-seq for H2B and H2Bub using a site-specific antibody that we recently developed (Van Welsem et al. 2018). First, we considered the patterns in the wild-type strain. Both the coverage at one representative locus and across all genes in a heatmap showed that H3K79me3 is predominantly present throughout coding sequences of most genes, where H2Bub is also high, as reported previously (Fig. 1D, 1E, Supplemental Fig. 1B; Schulze et al. 2009; Weiner et al. 2015; Sadeh et al. 2016; Magraner-Pardo et al. 2014). In contrast, H3K79me1 was found in transcribed as well as intergenic regions (Fig. 1D, Supplemental Fig. 1B). This is consistent with published ChIP-seq data and our previous ChIP-qPCR results (Weiner et al. 2015; Vlaming et al. 2016). In agreement with the distributive mechanism of methylation of Dot1 (Frederiks et al. 2008; De Vos et al. 2011), H3K79me1 and H3K79me3 anti-correlated, and H3K79me1 over the gene body was found on the minority of genes that lacked H3K79me3 and H2Bub (Fig. 1D). Among these low H3K79me3, high H3K79me1 genes were subtelomeric genes, where the Sir silencing complex competes with Dot1 for binding to nucleosomes and H2Bub levels are kept low by the deubiquitinating enzyme Ubp10 (Gardner et al. 2005; Emre et al. 2005; Gartenberg and Smith 2016; Kueng et al. 2013) (Supplemental Fig. 1C, 1E).

We then compared the patterns in wild-type versus *rpd3Δ* mutant strains. In metagene plots, the mutant showed a decrease in H3K79me1 and an increase in H3K79me3 just after the transcription start site (TSS; Supplemental Fig. 1B), suggesting that in this region Rpd3 suppresses the transition from lower to higher H3K79me states. To assess whether the changes observed in the metagene plots were explained by a modest effect on H3K79me at all genes or a stronger effect at a subset of genes, we determined the H3-normalized H3K79me3 level in the first 500 bp of each gene and ranked the genes based on the change in H3K79me3 upon loss of Rpd3. A heatmap of H3K79me3 by this ranking showed that the absence of Rpd3 leads to an increase of H3K79me3 at a subset of genes (Fig. 1F).

### Rpd3 represses H3K79me at its target genes

To characterize the genes at which H3K79me is regulated, we calculated the levels of H3K79me1 and H3K79me3 per gene in the same 500 bp window and plotted values in the rank order of H3K79me3 changes described above, using locally weighed regression (Fig. 2A; corresponding heatmaps can be found in Supplemental Fig. 2A). Inspection of these plots revealed that the ORFs on which H3K79me3 was increased in the *rpd3Δ* mutant showed a simultaneous decrease in H3K79me1 (groups III-IV; Fig. 2A). Strikingly, these Rpd3-regulated ORFs were on average marked with a relatively high level of H3K79me1 and low H3K79me3 in the wild-type strains but became more similar to the average yeast gene upon loss of Rpd3, consistent with the presence of a negative regulator of H3K79me acting on these ORFs. To test whether H3K79me-regulated genes were direct Rpd3 targets or indirectly affected, we assessed the relation between H3K79me changes and published data on Rpd3 binding and H4 acetylation (McKnight et al. 2015). The genes with the strongest increase in H3K79me3 upon Rpd3 loss had the highest Rpd3 occupancy, both at the promoter and in the 500 bp window downstream of the TSS (group IV; Fig. 2A). Rpd3 was also found to be active at these genes, since they were devoid of H4 acetylation in wild-type cells and H4 acetylation was restored in the *rpd3Δ* mutant (Fig. 2A). Finally, the top-regulated genes were also enriched for meiotic genes, which are known Rpd3 targets, and binding sites of Ume6, the Rpd3L subunit known to recruit Rpd3 to early meiotic genes (Fig. 2B, 2C) (Rundlett et al. 1998; Kadosh and Struhl 1998; Carrozza et al. 2005; Lardenois et al. 2015). Together, our results suggest that the genes at which Rpd3 restricts the build-up of H3K79me are direct targets of Rpd3.

**Figure 2.**
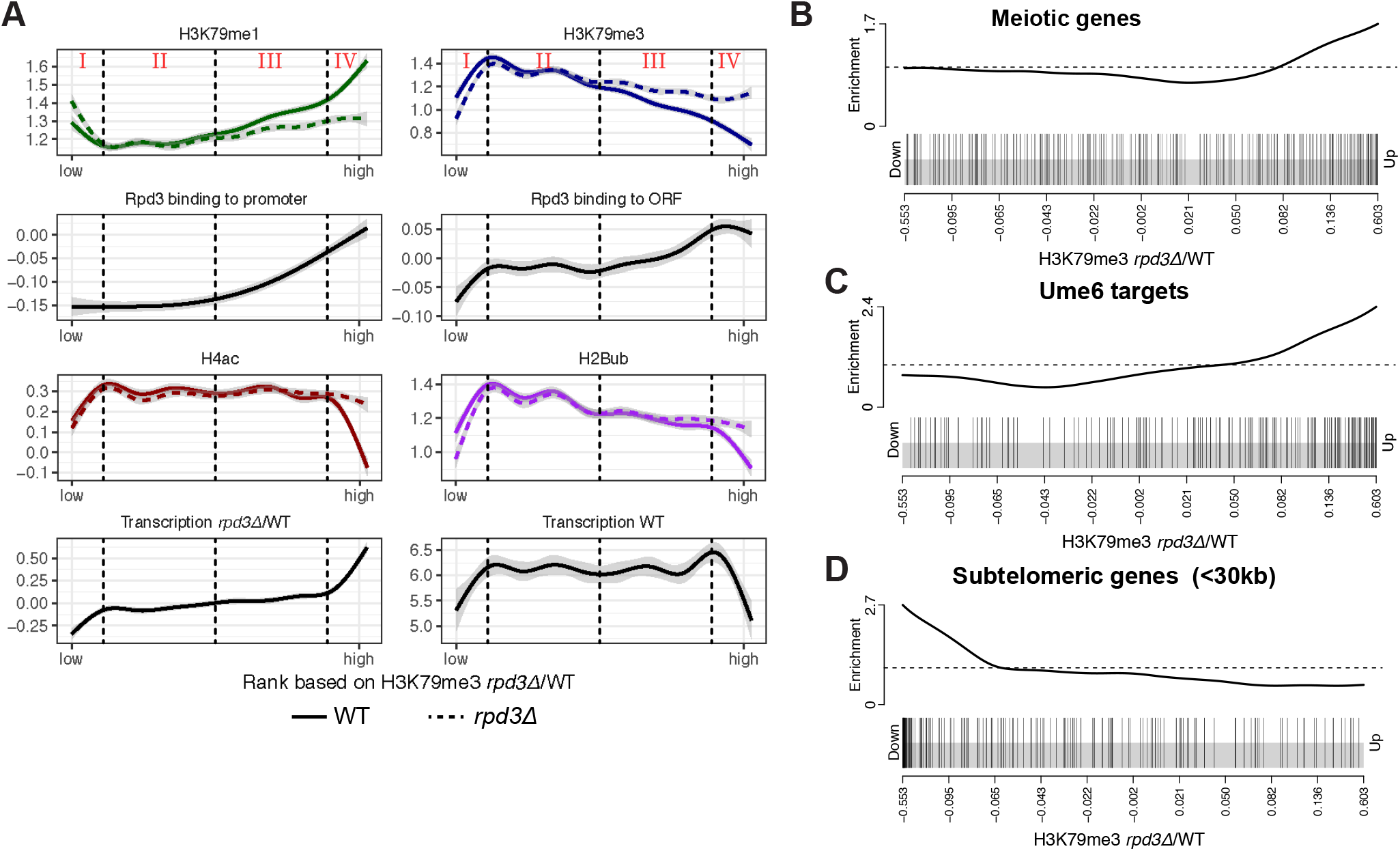
Rpd3 represses transcription, H2B ubiquitination and H3K79 methylation at its target sites. A) ChIP-seq and RNA-seq data for genes ranked on H3K79me3/H3 in *rpd3Δ/WT*, smoothed using locally weighed regression. The gray band around the line shows the 95% confidence interval. Vertical dashed lines separate 4 groups with distinct changes upon RPD3 deletion. For H3K79me1, H3K79me3, H2Bub, H4ac and Rpd3 binding to ORF, the average coverage in the first 500 bp was used. Rpd3 binding to promoter was the average coverage in the 400 bp upstream of the TSS. Transcription in wild-type cells was obtained from (McKnight et al. 2015), and the transcription in *rpd3Δ*/WT from (Kemmeren et al. 2014). B-D) Gene set enrichment analysis on genes ranked on H3K79me3/H3 in *rpd3Δ*/WT shows that meiotic (B) and Ume6-bound (C) genes are enriched among the genes at which Rpd3 represses H3K79 methylation, and subtelomeric genes (<30 kb of telomere) (D) are enriched among genes at which H3K79 methylation is decreased in *rpd3Δ* cells.

Notably, a small subset of genes loses H3K79me3 in the absence of Rpd3 (Fig. 1F, group I in Fig. 2A). This group of genes already has low H3K79me3 and H2Bub levels in wild-type cells and is highly enriched for subtelomeric genes (Fig. 2A, 2D). Loss of Rpd3 is known to enhance Sir-mediated silencing at subtelomeric regions (Ehrentraut et al. 2010, 2011; Gartenberg and Smith 2016; Thurtle-Schmidt et al. 2016). Our findings show that the stronger transcriptional silencing occurs with a concomitant reduction in H3K79me3 and H2Bub in the coding regions of heterochromatic genes. Whether or not the loss of these modifications contributes to the stronger silencing in *rpd3Δ/sin3Δ* mutants or is a consequence of it remains to be determined.

### Strong repression of H3K79me by Rpd3 coincides with repression of H2Bub and transcription

To understand the mechanistic basis for the crosstalk between Rpd3 and Dot1, we examined the changes in H2B ubiquitination and transcription. The expression of the H2Bub machinery is not deregulated in these mutants (Kemmeren et al. 2014) and no upregulation of H2Bub could be detected by immunoblot (Supplemental Fig. 1A). Because subtle changes can be missed by blot, we proceeded to generate H2Bub ChIP-seq data in wild-type and mutant strains using an antibody we recently described (Van Welsem et al. 2018). We found that the strongest H3K79me3 repression by Rpd3 (group IV) coincided with repression of H2Bub as well as transcription (Fig. 2A; RNA-seq data from (McKnight et al. 2015)). Moreover, these genes had below-average H2Bub and transcription levels in wild-type cells (Fig. 2A). H2Bub changes were confirmed by ChIP-qPCR (Supplemental Fig. 2B, 2C).

The H2B ubiquitination machinery is known to be recruited co-transcriptionally and promote H3K79 methylation, so transcriptional repression provides a likely explanation for the restriction of H3K79 methylation at these genes. However, despite these established causal links, there is no simple linear relation between transcription level and H3K79me3 level, while H2Bub correlates with transcription perfectly (Supplemental Fig. 2D; Schulze et al. 2009, 2011; Weiner et al. 2015). It appears that other processes counteract H3K79 methylation (see discussion), especially at highly transcribed genes, but that these processes do not affect the Rpd3-regulated genes as much, since they form a subset of genes at which transcription and H3K79me3, and their changes upon *RPD3* deletion, are correlated.

In addition to genes where Rpd3 has a strong effect on H3K79me, we also observed genes at which H3K79 methylation was more modestly affected by the deletion of *RPD3* (group III; Fig. 2A). Rpd3 is found at the promoters of these genes, but H4 acetylation, transcription, and H2B ubiquitination are not affected. Therefore, at these loci another, still unknown additional mechanism could be at play.

Taken together, we identified Rpd3 as a bona fide negative regulator of H3K79 methylation in yeast that restricts H3K79me3 at its euchromatic targets, probably mostly by repressing target gene transcription and H2Bub, but other mechanisms seem to be at play as well.

### HDAC1 loss increases H3K79me in murine thymocytes

Having uncovered a role for Rpd3 in restricting H3K79me at its targets and finding that this can explain a significant part of the H3K79 methylation variance between genes in yeast, we next wanted to investigate the biological relevance of this regulation in mammals. Histone deacetylases are conserved between species and can be divided into four classes (Yang and Seto 2008). Rpd3 is a founding member of the class I HDACs, which in mammals includes HDAC1, −2, −3 and −8. Of these, HDAC1 and - 2 are found in Sin3 complexes, like yeast Rpd3 (Yang and Seto 2008). Given that both HDACs and DOT1L play critical roles in T-cell malignancies, we employed conditional early thymocyte-specific *Hdac1* deletion (*Lck*-Cre;*Hdac1*^f/f^, resulting in *Hdac1^Δ/Δ^* thymocytes) in the mouse to investigate whether the regulation that we observed in yeast also exists in T cells. We focused on HDAC1 because it is more active in mouse thymocytes than HDAC2 (Heideman et al. 2013; Dovey et al. 2013). First, we measured the relative abundance of H3K79 methylation states on bulk histones by mass spectrometry in wild-type and *Hdac1*-deleted thymocytes of 3-week-old mice (Fig. 3A). In general, the overall levels of H3K79 methylation were much lower than in yeast and H3K79me1 was the most abundant methylation state, followed by H3K79me2, consistent with previous reports in mouse and human cells (Jones et al. 2008; Leroy et al. 2013). As seen in Figure 3A, *Hdac1*-deleted thymocytes had more H3K79me2 and H3K79me1 and less H3K79me0. Considering the distributive activity of Dot1 enzymes (Supplemental Fig. 3A), this suggests that Rpd3/HDAC1 is a conserved negative regulator of H3K79 methylation.

**Figure 3.**
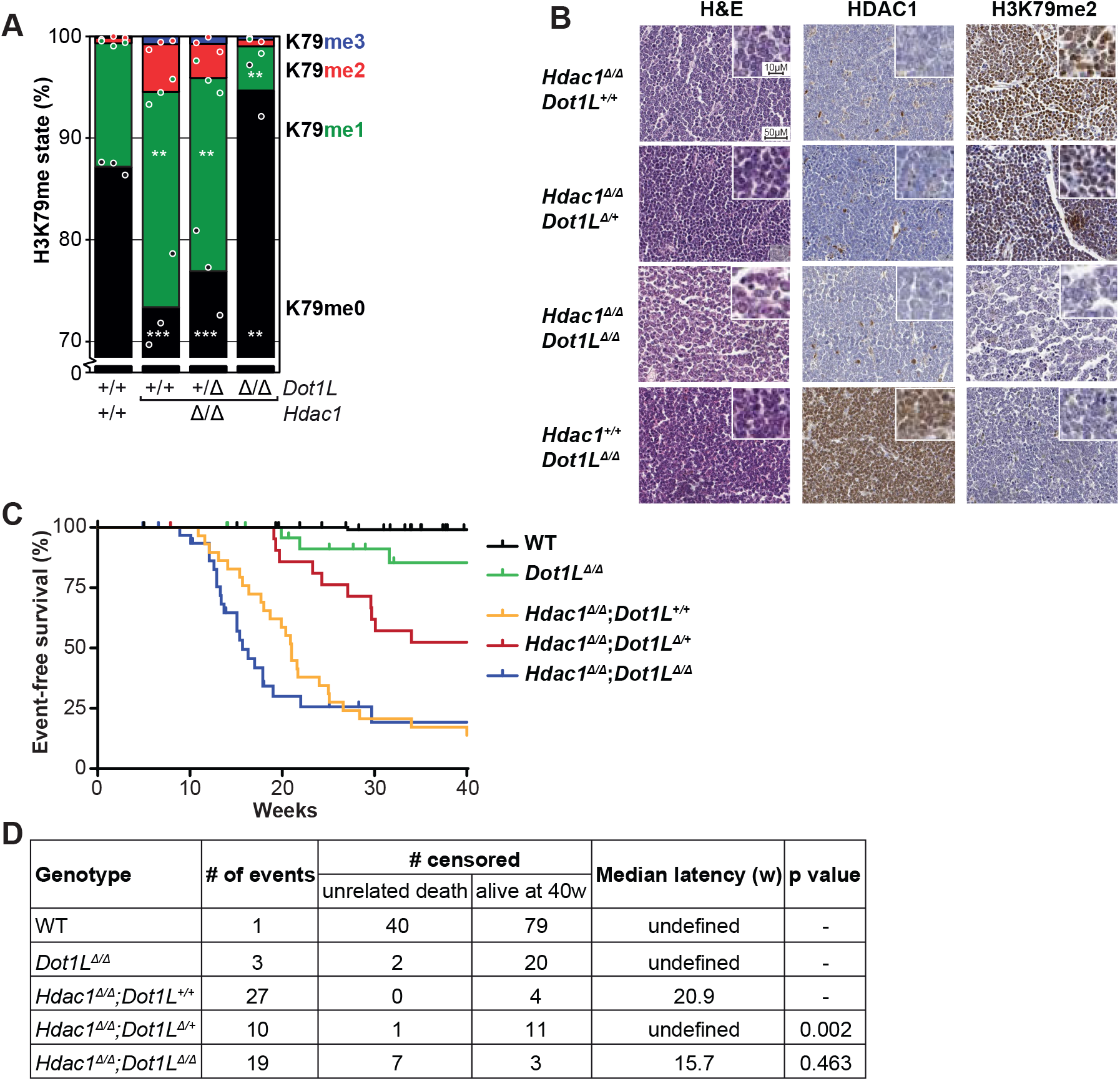
*Hdac1* deletion increases H3K79 methylation in thymocytes *in vivo,* and simultaneous heterozygous *Dot1L* deletion prolongs tumor-free survival. A) Mass spectrometry analysis of H3K79 methylation in thymuses from 3-week-old mice, either wild-type (Cre-) or with deleted *Hdac1* or *Dot1L* alleles, as indicated. Mean and individual data points of biological replicates; H3K79me0 is the predominant state, the y axis is truncated at 70% for readability. The remaining H3K79 methylation after homozygous *Dot1L* deletion is probably due to the presence of some cells in which Cre is not expressed (yet). B) Representative H&E and immunohistochemical stainings of thymic lymphomas of the indicated genotypes. A picture with lower magnification is included in Supplemental Fig. 3B. C) Kaplan-Meier curves of mice harboring thymocytes with indicated genotypes. An event was defined as death or sacrifice of a mouse caused by a thymic lymphoma. Mice that died due to other causes or were still alive and event-free at the end of the experiment were censored. Mice for which the cause of death could not be determined were removed from the data. Wild-type mice were the Cre-littermates of the mice that were used for the other curves. D) Summary of the data presented in panel C. A median latency could only be calculated when the tumor incidence was >50%. The p value was determined by comparing to the *Lck*-Cre;*Hdac1*^f/f^ curve with a Peto test, but a logrank test yielded the same conclusions.

### Reduced DOT1L dosage increases the latency of *Hdac1*-deficient thymic lymphomas

Conditional *Lck*-Cre;*Hdac1*^f/f^ knock-out mice die of thymic lymphomas characterized by loss of p53 activity and *Myc* amplification, with a 75% incidence and a 23-week mean latency (Heideman et al. 2013). Oncogenic transformation has not occurred yet in 3-week-old mice (Heideman et al. 2013), the age at which H3K79me levels were determined above. Since *Hdac1* deletion in thymocytes resulted in an increase in H3K79 methylation, as well as thymic lymphoma formation, we asked whether increased H3K79 methylation was important for tumor development in this mouse model. To address this question, a conditional *Dot1L* (*Dot1L*^f/f^) allele was crossed into the *Lck*-Cre;*Hdac1*^f/f^ line such that deletion of *Hdac1* was combined with deletion of zero, one or two *Dot1L* alleles. Immunohistochemistry confirmed the loss of HDAC1 at the protein level and mass spectrometry confirmed the loss of DOT1L protein activity for the expected genotypes (Fig. 3A, B, Supplemental Fig. 3B). H3K79me2 was used as an indicator for DOT1L presence, since none of the DOT1L antibodies we tested worked for IHC (antibody difficulties have also been described by Sabra et al. 2013).

Mice with conditional *Hdac1* alleles but wild-type for *Dot1L* (*Lck*-Cre;*Hdac1*^f/f^) developed thymic lymphomas for which they had to be sacrificed, with a median latency of 21 weeks and an incidence of 86% during the 40-week length of the experiment, comparable to what was observed before (Fig. 3C, D) (Heideman et al. 2013). As expected, Lck-Cre-negative control mice rarely developed thymic lymphomas (1 out of 112). Also, *Dot1L* deletion alone (*Lck*-Cre;*Dot1L*^f/f^) rarely led to thymic lymphomas, with a 15% incidence in this background (Fig. 3C, 3D), and no cases of thymic lymphoma in another background (data not shown). We then assessed the effect of *Dot1L* deletion in the *Lck*-Cre;*Hdac1*^f/f^ model. Loss of one copy of *Dot1L* increased survival rate and tumor latency (48% incidence, comparison to *Hdac1*^f/f^ alone p=0.002; Fig. 3C, D). This effect suggests that there is a causal link between the increase in H3K79 methylation and the development or maintenance of thymic lymphomas upon *Hdac1* deletion. Interestingly, homozygous *Dot1L* deletion, leading to a complete loss of H3K79me, did not extend the latency of thymic lymphomas (81% incidence, 15.7 weeks median latency, comparison to *Hdac1* deletion alone p=0.463; Fig. 3C, D). A possible explanation is that the simultaneous deletion of *Dot1L* and *Hdac1* results in the generation of a different class of tumor that does not depend on H3K79 methylation but has acquired other, possibly epigenetic, events that allow oncogenic transformation. A similar model has been proposed for the loss of *Hdac2* in the *Lck*-Cre;*Hdac1*^f/f^ model (Heideman et al. 2013). For the heterozygous *Dot1L* effect, we consider two possible explanations: the oncogenic transformation occurred later because an additional event was required to overcome the lack of high H3K79me, or tumors grew slower due to lower H3K79me, either because of decreased proliferation or increased apoptosis.

### *Hdac1*-deficient thymic lymphoma lines depend on DOT1L activity

To further study the *Dot1L* dependence of *Hdac1*-deficient thymic lymphomas in a more controlled environment, we turned to *ex vivo* experiments. Cell lines were derived from *Hdac1*-deficient thymic lymphomas (Heideman et al. 2013). Since these cell lines had an inactivating mutation in p53, cell lines derived from p53-null thymic lymphomas were used as *Hdac1*-proficient control lines (Heideman et al. 2013). First, we examined whether *Hdac1*-deficient tumor cells retained the increased H3K79 methylation levels seen prior to the oncogenic transformation. Both by immunoblot and by targeted mass spectrometry on independent samples (Fig. 4A, 4B), *Hdac1*-deficient tumor cell lines had more H3K79 methylation than their *Hdac1*-proficient counterparts. Thus, the effect of HDAC1 on H3K79 methylation observed *in vivo* in 3-week old pre-malignant thymuses was maintained in the thymic lymphoma cell lines. Importantly, *Hdac1*-deficient cell lines also possessed high levels of ubiquitinated H2B compared to *Hdac1*-proficient controls (Fig. 4C). High H2Bub is consistent with the increase in H2Bub seen at Rpd3 targets in yeast. Therefore, a plausible model is that at least part of the observed increase in H3K79me upon loss of HDAC1 activity is mediated via H2Bub, consistent with what we observed in yeast. However, contributions from other regulatory mechanisms cannot be excluded (see discussion). To test the DOT1L dependence of the cell lines, DOT1L was depleted using shRNAs in *Hdac1*-proficient and -deficient cell lines. As can be seen in Figure 4D, shRNAs that reduce Dot1L expression (Supplemental Fig. 4A) affected proliferation of the *Hdac1*-deficient cell lines. Compared to the control lines, the *Hdac1*-deficient cell lines were also more sensitive to two different DOT1L inhibitors (Fig. 4E). Both inhibitors, EPZ-5676 (Pinometostat) and SGC-0946, effectively lowered H3K79 methylation (Supplemental Figure 4B). Thus, shRNA-mediated DOT1L knockdown and chemical DOT1L inhibition showed that the *Hdac1*-deficient thymic lymphoma cell lines depended on DOT1L activity. The reduced growth upon inactivation of DOT1L could be explained by a block in proliferation or an increase in cell death. To measure apoptosis induction, the levels of AnnexinV and DAPI staining of non-permeabilized cells were determined by flow cytometry. In the DMSO-treated condition most cells were alive, although *Hdac1*-deficient lines had a slightly higher basal apoptosis level (Fig. 5A,B). This combination of proliferation and apoptosis has also been observed in *Hdac1^Δ/Δ^* teratomas (Lagger et al. 2010). However, DOT1L inhibition by 5μM of SGC-0946 dramatically induced apoptosis in *Hdac1*-deficient cells, whereas no effect on apoptosis was observed in the control cell lines (Fig. 5A,B). Thus, DOT1L inhibition induced apoptosis specifically in *Hdac1*-deficient thymic lymphoma cell lines.

**Figure 4.**
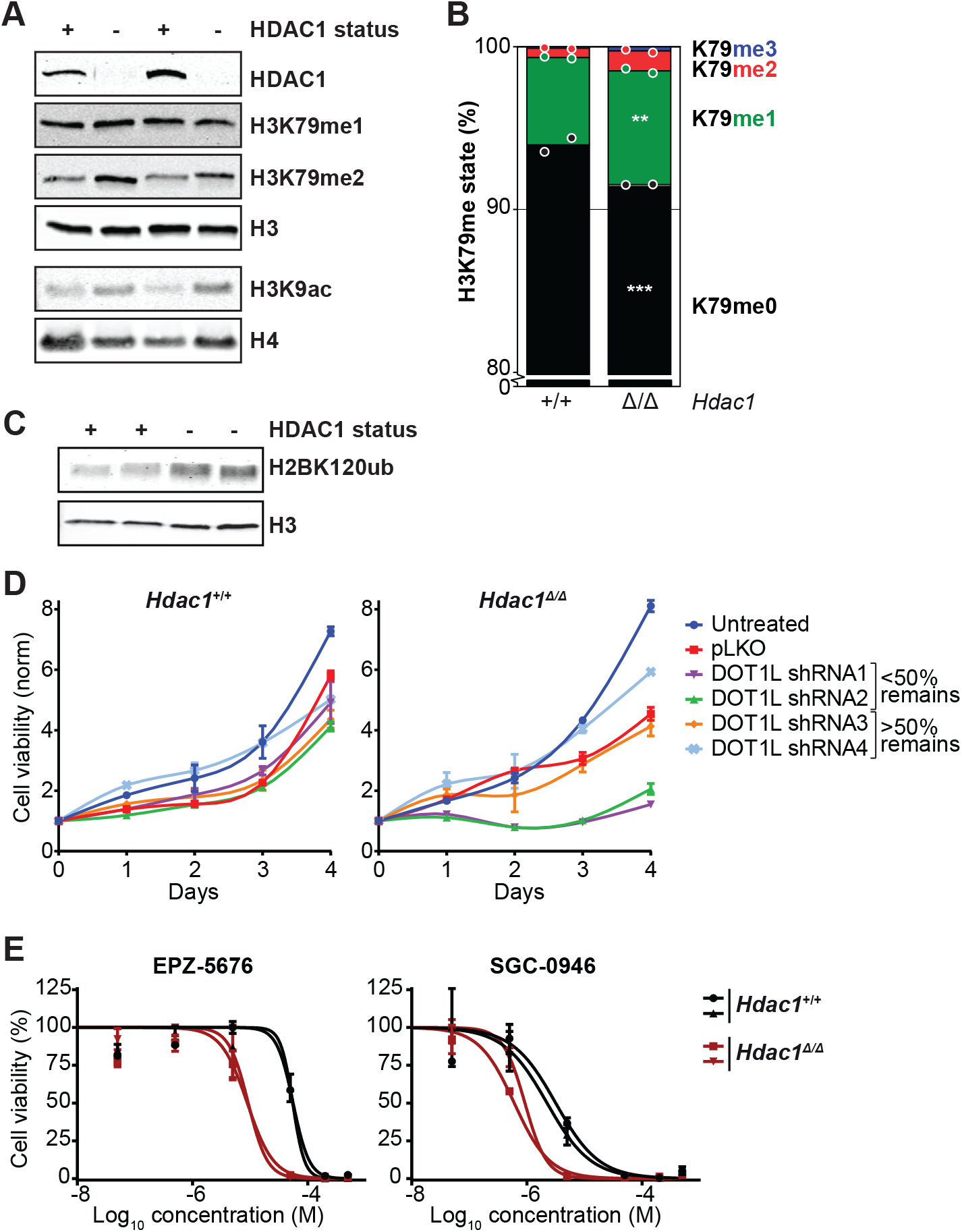
*Hdac1*-deleted thymic lymphoma cell lines depend on DOT1L activity. A) Immunoblots showing HDAC1 status and H3K79me/H3K9ac levels in nuclear lysates of *Hdac1*-proficient (p53-null) and *Hdac1*-deficient thymic lymphoma cell lines. The top four and bottom two panels are from separate lysates of the same cell lines. B) Mass spectrometry analysis of H3K79 methylation in the cell lines from panel A. Mean and individual data points of two independent cell lines; H3K79me0 is the predominant state, the y axis is truncated at 80% for readability. C) Immunoblot showing H2BK120 ubiquitination levels in *Hdac1*-proficient and -deficient cell lines (two independent lines each). D) Growth curves of *Hdac1*-proficient and -deficient cell lines that were left untreated or were infected with empty virus (pLKO) or shRNAs against *Dot1L* and selected with puromycin. Growth curves were determined by a series of resazurin assays, which measure metabolic activity, starting four days post-infection. Error bars indicate the range of two replicates from independent cell lines. E) Inhibitor dose-response curves of the two DOT1L inhibitors EPZ-5676 (Pinometostat) and SGC-0946 in *Hdac1*-proficient and -deficient cell lines. Cell viability was measured by a resazurin assay after three days of treatment and measurements were normalized to DMSO-treated cells. Two independent cell lines are plotted separately; error bars represent the range of two biological replicates.

**Figure 5.**
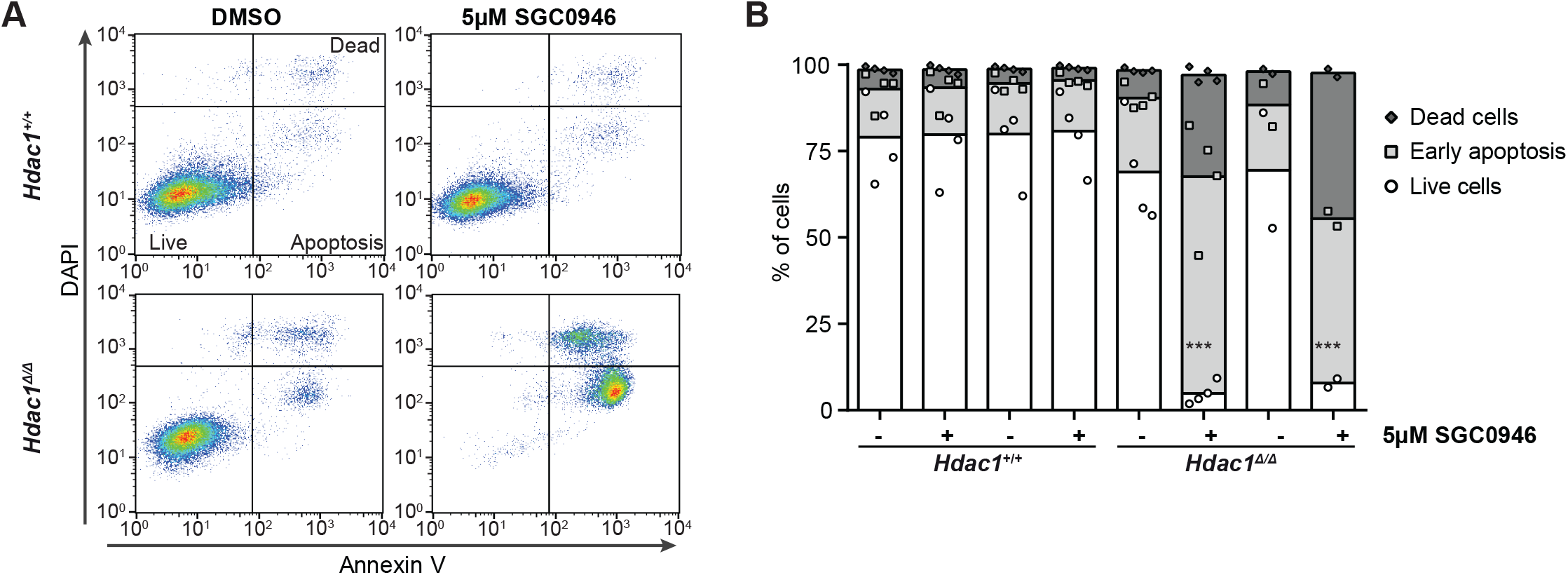
*Hdac1*-deficient thymic lymphoma lines require DOT1L activity for survival. A) Representative apoptosis FACS plots of cell lines treated with DMSO or the DOT1L inhibitor SGC-0946 for two days. Annexin V staining and DAPI staining were performed on unpermeabilized cells to distinguish live (Annexin V low; DAPI low), apoptotic (Annexin V high; DAPI low) and dead (Annexin V high; DAPI high) cells. B) Quantification of several independent FACS experiments, including the experiment shown in panel A. Mean (bars) with individual data points of 2-4 replicates each of two independent lines per genotype.

## Discussion

Here we describe that the yeast HDAC, Rpd3, is a negative regulator of H3K79 methylation that restricts methylation at the 5’ ends of its target genes. Similar to what we observe for Rpd3 in yeast, deleting *Hdac1* in murine thymocytes leads to an increase in H3K79 methylation. This regulation is relevant in a cancer context, since heterozygous deletion of *Dot1L* prolongs the survival of mice that develop *Hdac1*-deficient thymic lymphomas. Cell lines derived from *Hdac1*-deficient thymic lymphomas undergo apoptosis upon DOT1L inhibition or depletion, which indicates a form of non-oncogene addiction to DOT1L.

### Rpd3 target genes

In the yeast genome, most euchromatic genes are marked by H2Bub and H3K79me3 in their transcribed region. While the levels of H2Bub correlate well with transcription levels, it is evident that H2Bub is not the only determinant of H3K79me3 in yeast because the relation between transcription and H3K79me3 is more complex (Fig. 2A). While genes silenced by the SIR complex have low H3K79me3 levels due to active repression mechanisms (Gartenberg and Smith 2016), the majority of euchromatic genes contain H3K79me3 irrespective of their expression level (Supplemental Fig. 2D; Schulze et al. 2009, 2011; Weiner et al. 2015). Some genes contain lower H3K79me3 and higher H3K79me1 levels than the average gene, however. A subset of these deviants has been identified as genes undergoing antisense transcription (Murray et al. 2015; Brown et al. 2018), possibly resulting in nucleosome instability and increased histone turnover, which counteracts the buildup of higher H3K79me states but does not affect the more dynamic H2Bub modification (Weiner et al. 2015). Here we provide insight into the low H3K79me3/high H3K79me1 levels of another subset of yeast genes. H3K79me ChIP-seq in yeast revealed that Rpd3 restricts H3K79me3 at its direct target genes. This subset of genes showed great overlap with the minority of euchromatic genes that is marked with H3K79me1 instead of H3K79me3. Thus, the regulation by Rpd3 provides an explanation for the variation in H3K79 methylation between genes and thereby seems to be an important determinant of the H3K79 methylation pattern. The H3K79me effect of Rpd3 was most notable at the 5’ end of genes, which is in agreement with previous studies on Rpd3 activity. The deacetylation activity of Rpd3/Rpd3L is reported to be strongest in coding sequences, particularly at the 5’ ends (Weinberger et al. 2012). At which genes HDAC1 regulates H3K79 methylation in murine thymocytes is an interesting question, but addressing it is not straight-forward. Unlike in yeast, in mammals H3K79 methylation is tightly linked to the transcriptional activity at genes. Processes through which transcription promotes H3K79 methylation are known, but in turn, H3K79 methylation may affect transcription as well (reviewed in Vlaming and Van Leeuwen 2016). Therefore, while assessing H3K79me changes in *Hdac1*-deficient cells, it will be challenging to separate direct effects from indirect effects on DOT1L activity through transcriptional changes.

### Mechanism of regulation

What could be the mechanism of H3K79me regulation by Rpd3/HDAC1? Until now, the H2B ubiquitination machinery was the only described H3K79me regulator conserved from yeast to mammals (Weake and Workman 2008). Here we describe another conserved regulator, Rpd3/HDAC1, and our results indicate that it acts in part by restricting H2Bub. Although transcription and H3K79 methylation are not clearly positively correlated in wild-type yeast cells (Supplemental Fig. 2D; Weiner et al. 2015; Magraner-Pardo et al. 2014; Schulze et al. 2011), we observed that the H3K79 methylation changes in *rpd3Δ* correlate very well with transcriptional changes. These data, together with the well-established causal relationships between transcription and H2B ubiquitination on the one hand and H2B ubiquitination and H3K79 methylation on the other hand, suggest that there is indeed a causal link between (sense) transcription and the placement of H3K79 methylation at a subset of the yeast genome. At higher transcription levels however, this relationship can be obscured by other processes, most likely histone turnover, counteracting the high Dot1 activity (Radman-Livaja et al. 2011; Murray et al. 2015). We have recently identified the conserved histone acetyltransferase Gcn5 as a negative regulator of H3K79me and H2Bub (Vlaming et al. 2016). At first glance, it seems counterintuitive that a HAT and an HDAC have overlapping effects. However, acetylation at non-overlapping histone or non-histone lysines may explain this discrepancy. For example, Gcn5 most likely negatively regulates H2Bub and H3K79me by affecting the deubiquitinating module of the SAGA co-activator complex in which Gcn5 also resides (Vlaming et al. 2016). H2B ubiquitination by human RNF20/40 has also been shown to be regulated by histone acetylation (Garrido Castro et al. 2018). Treatment of acute lymphoblastic leukemia cell lines with the non-selective HDAC inhibitor Panobinostat showed changes in H2Bub, with decreased H2Bub in MLL-r leukemia lines and increased H2Bub in non MLLr-leukemia lines, suggesting context-dependent mechanisms (Garrido Castro et al. 2018).

Besides H2Bub, other mechanisms are likely to contribute to the observed H3K79me increase in the absence of Rpd3/HDAC1 as well. Histone acetylation is increased in the absence of the deacetylase HDAC1, and histone acetylation has been previously linked to DOT1L recruitment through the YEATS domain transcription elongation proteins AF9 and ENL (Li et al. 2014; Kuntimaddi et al. 2015; Erb et al. 2017; Wan et al. 2017). Very recently, preferential Dot1 binding to acetylated H4K16 has been shown, and the histone acetyltransferase Sas2 was found to be a positive regulator of H3K79 methylation in yeast, probably via acetylation of H4K16 (Lee et al. 2018). Our identification of Rpd3/HDAC1 as a regulator of DOT1L underscores the intimate relationship between histone acetylation and H2Bub and H3K79me and provides evidence for a specific HDAC involved in the crosstalk: HDAC1.

### DOT1L in tumor maintenance

Loss of HDAC1 leads to oncogenic transformation and higher H3K79me in murine thymocytes. Heterozygous *Dot1L* deletion prolonged the survival of mice with thymocyte-specific *Hdac1* deletion due to a lower incidence and increased latency of thymic lymphomas. Our analysis of *Hdac1*-deficient thymic lymphoma cells *ex vivo* provided more insight into the possible mechanisms for the reduced tumor burden. Using DOT1L inhibitors and a knock-down approach, we established that DOT1L was required for survival of the tumor cells by preventing the induction of apoptosis, suggesting that DOT1L is required for tumor maintenance. The DOT1L dependency of the thymic lymphomas resembles that of MLL-rearranged leukemia (Wang et al. 2016) as well as breast and lung cancer cell lines (Kim et al. 2012; Zhang et al. 2014). The full genetic deletion of *Dot1L* did not reduce tumor burden. The reasons for this are currently unknown and require further study. One possible reason is that some remaining DOT1L activity and H3K79 methylation might be required to induce apoptosis in the tumor cells. We note that in the *ex vivo* experiments where *Dot1L* knock down and DOT1L inhibitors were found to lead to induction of apoptosis, some residual H3K79me was indeed still present. Another possibility is that the simultaneous loss of *Hdac1* and *Dot1L* imposes oncogenic transformation through alternative, epigenetic mechanisms that bypass the apoptotic-prone state. This would be in agreement with the known role of DOT1L in the maintenance of cellular epigenetic states (Onder et al. 2012; Soria-Valles et al. 2015; Breindel et al. 2017). Regardless of possible mechanisms, the finding that HDAC1 affects DOT1L activity in yeast as well as mouse T cells, warrants further investigation. For example, it will be interesting to determine whether and under which conditions HDAC1 activity influences DOT1L activity in human MLL-r leukemia and whether the crosstalk is involved in the response of CTLC to HDAC inhibitors in the clinic. Our findings in murine lymphoma add to a growing list of cancers that rely on DOT1L activity, and therefore underline the importance of understanding the regulation of DOT1L.

## Materials and methods

### Yeast strains

Yeast strains used in this article are listed in Supplemental Table 1. Yeast media were described previously (Van Leeuwen and Gottschling 2002). The generation of the barcoded deletion library used for the Epi-ID experiment was described previously (Vlaming et al. 2016). Yeast *rpd3Δ* and *sin3Δ* strains were taken from this library and independent clones were generated by deleting these genes in the barcoded wild-type strain NKI4657, using the NatMX selection marker from pFvL99 (Stulemeijer et al. 2011). Gene deletions were confirmed by PCR.

### Cell culture

Thymic lymphoma cell lines (Heideman et al. 2013) were cultured under standard conditions in RPMI 1640 (Gibco) supplemented with 10% FBS (Sigma-Aldrich), antibiotics and L-Glutamine. HEK 293T cells were cultured in DMEM (Gibco) supplemented with 10% FBS (Sigma-Aldrich) and L-Glutamine.

### Mouse survival analysis

The generation and crosses of the conditional knock-out mice is described in the Supplemental Methods. Mice were monitored over time, up to 40 weeks of age. A power analysis performed beforehand determined group sizes of 20-25 mice per genotype. Mice were sacrificed before 40 weeks when they displayed breathing issues caused by the thymic lymphoma, or serious discomfort unrelated to tumor formation. After death or sacrifice, mice were checked for the presence of a thymic lymphoma. Mice that died without a lymphoma were censored in the survival curves. Mice of which the cause of death could not be determined were left out of the survival curve, together with their littermates. All experiments were approved by a local ethical committee and performed according to national guidelines. Mice were housed under standard conditions of feeding, light and temperature with free access to food and water.

### Protein lysates, immunoblots and mass spectrometry

Yeast whole-cell extracts were made as described previously (Vlaming et al. 2014). Nuclear extracts were prepared of murine cells, see supplemental methods for details. The immunoblotting procedure was as described in (Vlaming et al. 2014). All antibodies used in this study are listed in the supplemental methods. Mass spectrometry measurements on yeast strains and thymic lymphoma cell lines were as described in (Vlaming et al. 2014). Measurements on thymus tissue were performed using the method described in (Vlaming et al. 2016).

### ChIP-sequencing and ChIP-qPCR

ChIP and ChIP-qPCR experiments were performed as described before (Vlaming et al. 2016). Primers for qPCR are shown in Supplemental Table 2. Details on the ChIP-seq library preparation and the first analysis steps are provided in the Supplemental Methods. In short, preparation and sequencing were performed by the NKI Genomics Core Facility. Reads were mapped to the *Saccharomyces cerevisiae* reference genome R64-2-1 and extended to 150 bp. Samples were depth-normalized, and when data from biological duplicates was found to be similar, data sets were merged for further analyses. Metagene plots and heatmaps were generated with custom scripts in R/Bioconductor (Cherry et al. 2012; Huber et al. 2015). Reads were aligned in a window of −500 to 1 kb around the TSS of each verified ORF recorded in SGD. Genes that contained a coverage of 0 or an average coverage in the first 500 bp below 0.5 were filtered out (leaving 5006 out of 5134 genes). For heatmaps, the coverage was grouped in bins of 10 bp. The plots in Figure 2A and Supplemental Fig. 2D were created with custom scripts in R by using locally weighed regression (LOESS). Transcription in WT cells was obtained from (McKnight et al. 2015), and transcription in *rpd3Δ*/WT from (Kemmeren et al. 2014). Gene set enrichment plots were created with the barcodeplot function from the limma package (Ritchie et al. 2015). Ume6 targets were obtained from the Yeastract database (Teixeira et al. 2014) and filtered for simultaneous DNA binding and expression evidence. The list of meiosis factors was generated by searching for genes with a GO term containing “meio” or children thereof. The distance of each gene from the telomere on the same chromosome arm was calculated manually by using genome feature information from SGD.

### Histology/immunohistochemistry

Tissues were fixed in EAF (ethanol, acetic acid, formol saline) for 24 hours and subsequently embedded in paraffin. Slides were stained with hematoxylin and eosin (H&E), or immunohistochemistry was performed as described (Heideman et al. 2013).

### Knockdown/viability assays and Dot1L mRNA analysis

To produce lentiviral particles, HEK 293T cells were co-transfected with shRNA-containing pLKO.1 and three packaging plasmids containing Gag and Pol, Rev and VSV-G, respectively, using PEI. The media was replaced after 16 hours and virus-containing medium was harvested 72 hours after transfection. Virus particles were 10x concentrated from filtered medium using AmiCon 100 kDa spin columns. For lentiviral infections, 100,000 cells were seeded in 96-well tissue culture plates and infected using 7.5 μL concentrated virus, in the presence of 8 μg/ml Polybrene. The medium of infected cells was replaced with puromycin-containing medium 48 hours after infection and refreshed again 72 hours after infection, after which a cell viability assay was performed every 24 hours. Cell viability was determined by a CellTiter-Blue (Promega) assay, measuring conversion to resorufin after three hours with the EnVision Multilabel Reader (PerkinElmer). All cells treated with a particular shRNA were pooled for RNA isolation using the RNeasy Mini kit (Qiagen). DNaseI (New England Biolabs) digestion was performed and RNA was reverse transcribed into cDNA using SuperScript II Reverse Transcriptase (Invitrogen) according to the manufacturer’s protocol. *Dot1L* transcript abundance was measured by qPCR using SYBRgreen master mix (Roche) and the LightCycler 480 II (Roche). qPCR primers can be found in Supplemental Table 2.

### Inhibitor treatment

100,000 cells were seeded in 96-well tissue culture plates, in 200 μL culture medium containing the indicated concentration of inhibitor. Two inhibitors were used: SGC-0946 (Structural Genomics Consortium) and EPZ-5676 (Pinometostat; Selleck Chemicals). Cell viability was determined after three days, as described above. Data was normalized, with the maximum of each cell line to 100% and background fluorescence set to 0%. Graphpad Prism was used to fit log(inhibitor) vs normalized response curves with a variable slope.

### Apoptosis FACS

Cells treated with 0.1% DMSO or 5 μM SGC-0946 were stained with Annexin V-FITC and DAPI following the protocol of the Annexin V-FITC Apoptosis Detection Kit (Abcam). Fluorescence was detected by FACS using the CyAn ADP Analyzer (Beckman Coulter) and data was analyzed using FlowJo software.

### Statistics

Survival curves were plotted in Graphpad Prism, and Peto Mortality-Prevalence tests were performed in SAS to compare the curve of Lck-Cre;Hdac1^f/f^ mice with the *Lck*-Cre;*Hdac1*^f/f^;*Dot1L*^f/WT^ and *Lck*-Cre;*Hdac1*^f/f^;*Dot1L*^f/f^ curves. The same conclusions could be drawn based on the standard logrank test in Graphpad Prism. Mass spectrometry data were compared using two-way ANOVA, comparing samples of all genotypes to the wild-type sample and using the Dunnett correction for multiple comparisons. The significance thresholds used were p<0.05 (*), p<0.01 (**) and p<0.001 (***). The asterisks indicate significant differences compared to wild type.

### Accession numbers

The accession number for the ChIP-seq results reported in this study is GSE107331.

## Acknowledgements

The authors thank Jeffrey McKnight for sharing Rpd3 and H4ac ChIP-seq data. We thank Roel Wilting and Marinus (Richard) Heideman for help with initial HDAC1 experiments and Michael Hauptman for statistical tests on the survival curves. We thank the NKI animal pathology facility for histology and immunohistochemistry, as well as advice, the NKI genomics core facility for library preparations and sequencing, the NKI FACS facility for assistance, Onno Bleijerveld for mass spectrometry advice, and the caretakers of the NKI lab animal facility for assistance and excellent animal care. We thank Ila van Kruijsbergen, Tineke Lenstra, and Maarten van Lohuizen for critically reading the manuscript.

This work was supported by the Dutch Cancer Society (KWF2009-4511 and NKI2014-7232 to FvL and HJ) and the Netherlands Organisation for Scientific Research (NWO-VICI-016.130.627, NCI-KIEM-731.013.102 and NCI-LIFT-731.015.405 to FvL and ZonMW Top 91213018 to HJ). The funders had no role in study design, data collection and interpretation, or the decision to submit the work for publication.

HV, CMM and FvL designed the studies and the cell line and mouse studies together with HJ and JHD. Yeast experiments were performed by HV, TM and TvW, mouse experiments by CMM, HV, SH, EMK-M, MFA and CL, and cell line experiments by CMM, SP, and HV. TK performed ChIP-seq and genomics analyses, SK made IHC pictures and gave histology advice, LH and TTS performed mass spectrometry measurements and were advised by AFMA. HV and FvL wrote the manuscript, with help from CMM, HJ and JHD.

The Netherlands Cancer Institute and FvL are entitled to royalties that may result from licensing the yeast H2BK123ub-specific monocloncal antibody according to IP policies of the Netherlands Cancer Institute. The other authors declare that no competing interests exist.

## Supplemental Material

**Supplemental Table 1.** Yeast strains used in this study

| Strain name | Genotype | Reference | Figures |
| --- | --- | --- | --- |
| NatMX-KO x Barcoders library | MATa can1Δ::STE2pr-Sphis5 lyp1Δ his3Δ1 leu2Δ0 ura3Δ0 met15Δ0 hoΔ::barcodedKanMX GOI::NatMX | (Vlaming et al. 2016) | 1B |
| NKI4560 | MATa can1Δ::STE2pr-Sp_his5 lyp1Δ his3Δ1 leu2Δ0 ura3Δ0 met15Δ0 hoΔ::barcode(0001)KanMX | (Vlaming et al. 2016) | 1C, S1A |
| NKI4557 | MATa can1Δ::STE2pr-Sp_his5 lyp1Δ his3Δ1 leu2Δ0 ura3Δ0 met15Δ0 hoΔ::barcode(sir2)KanMX dot1Δ::NatMX | (Vlaming et al. 2016) | S1A |
| NKI4558 | MATa can1Δ::STE2pr-Sp_his5 lyp1Δ his3Δ1 leu2Δ0 ura3Δ0 met15Δ0 hoΔ::barcode(sir3)KanMX bre1Δ::NatMX | (Vlaming et al. 2016) | S1A |
| NKI4643 | MATa can1Δ::STE2pr-Sp_his5 lyp1Δ his3Δ1 leu2Δ0 ura3Δ0 met15Δ0 hoΔ::barcode(0791)KanMX rpd3Δ::NatMX | From library | 1C, S1A |
| NKI4644 | MATa can1Δ::STE2pr-Sp_his5 lyp1Δ his3Δ1 leu2Δ0 ura3Δ0 met15Δ0 hoΔ::barcode(1002)KanMX sin3Δ::NatMX | From library | 1C, S1A |
| NKI4657 | MATa his3Δ200 leu2Δ0 trp1Δ63 ura3Δ0 met15Δ0 hoΔ::barcode(sir4)KanMX | (Vlaming et al. 2016) | All ChIPs |
| NKI4713 | MATa his3Δ200 leu2Δ0 trp1Δ63 ura3Δ0 met15Δ0 hoΔ::barcode(sir4)KanMX rpd3Δ::NatMX | This study | All ChIPs |

**Supplemental Table 2.**
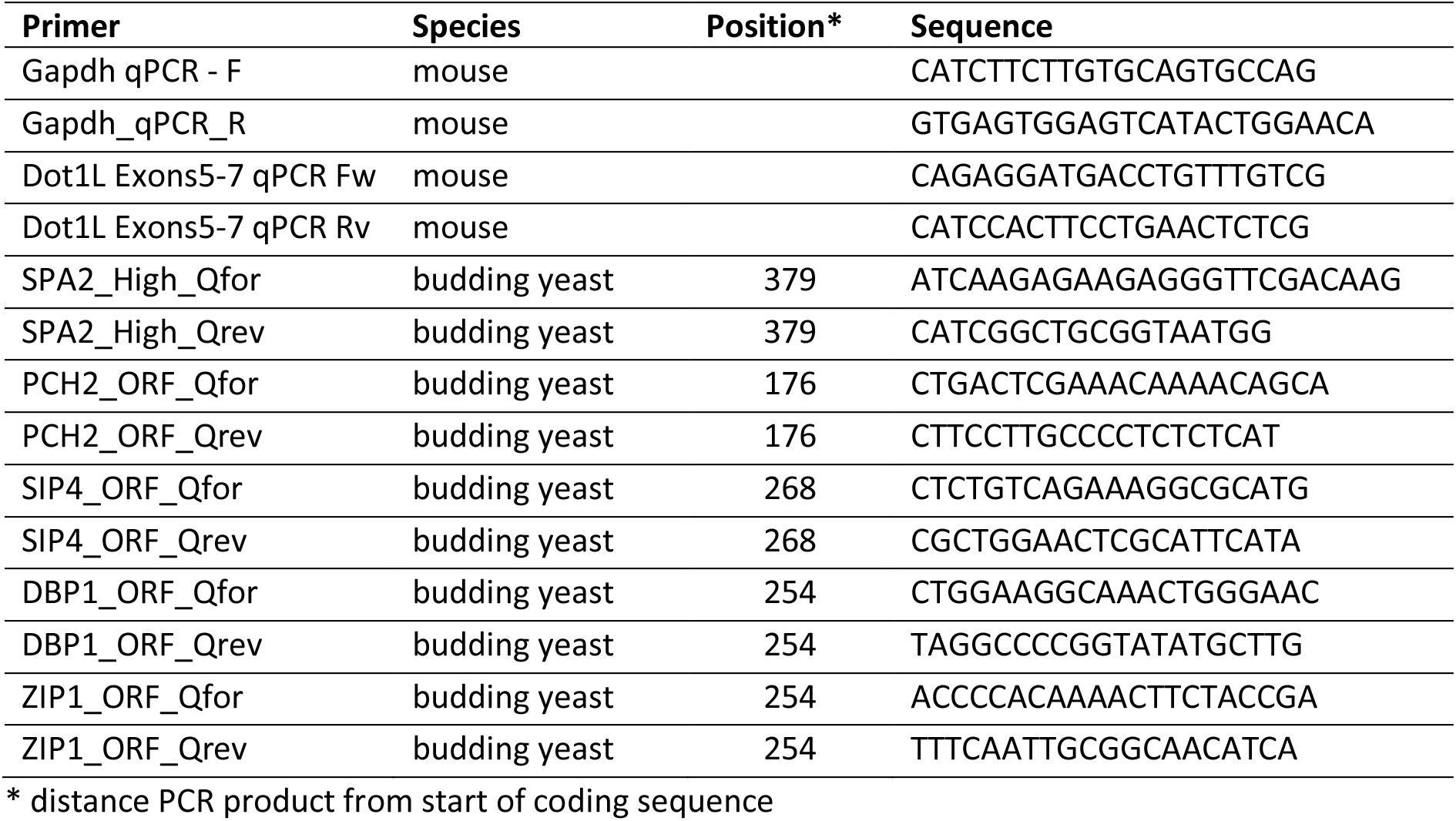
Primers used for qPCR

**Supplemental Figure 1.**
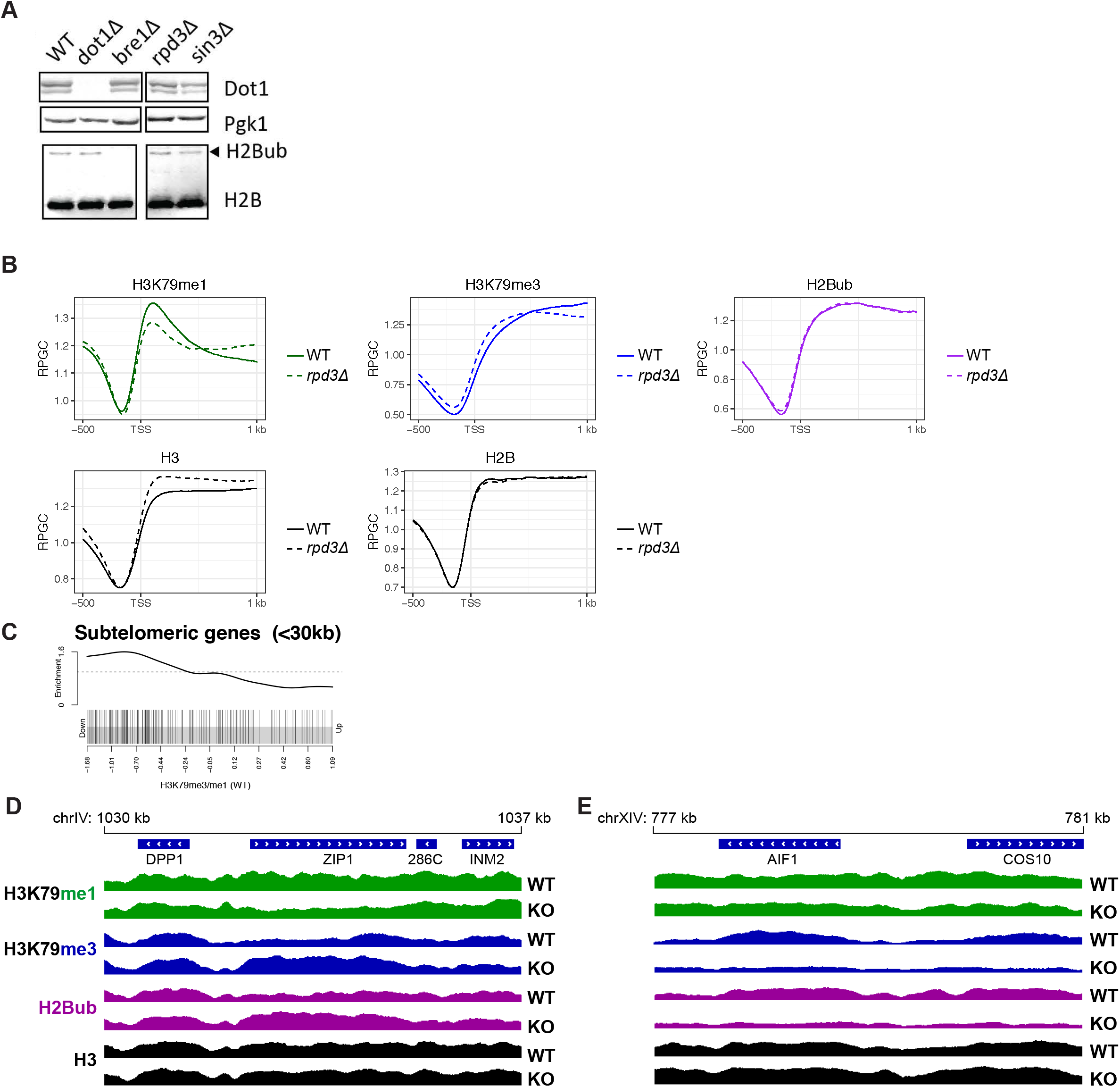
Rpd3 and Sin3 negatively regulate H3K79 methylation. A) Immunoblots show that deletion of *RPD3* or *SIN3* does not lead to a detectable increase in global H2Bub or Dot1 protein levels. B) Metagene plots of H3K79me1, H3K79me3, H2Bub, H3, and H2B in *rpd3Δ* and wild-type strains. C) Gene set enrichment analysis shows that subtelomeric genes (<30 kb of telomeres) are enriched among genes with low H3K79 methylation (measured by the average H3K79me3/H3K79me1 ratio in the first 500 bp of the ORF). D-E) Snapshots of depth-normalized ChIP-seq data tracks from wild-type and *rpd3Δ* strains showing 7 kb surrounding the meiotic gene *ZIP1* (D), and telomeric genes *AIF1* and *COS10* (E). All tracks have the same y axis (0-2), which, for comparison, is also the same scale as in Fig. 1E.

**Supplemental Figure 2.**
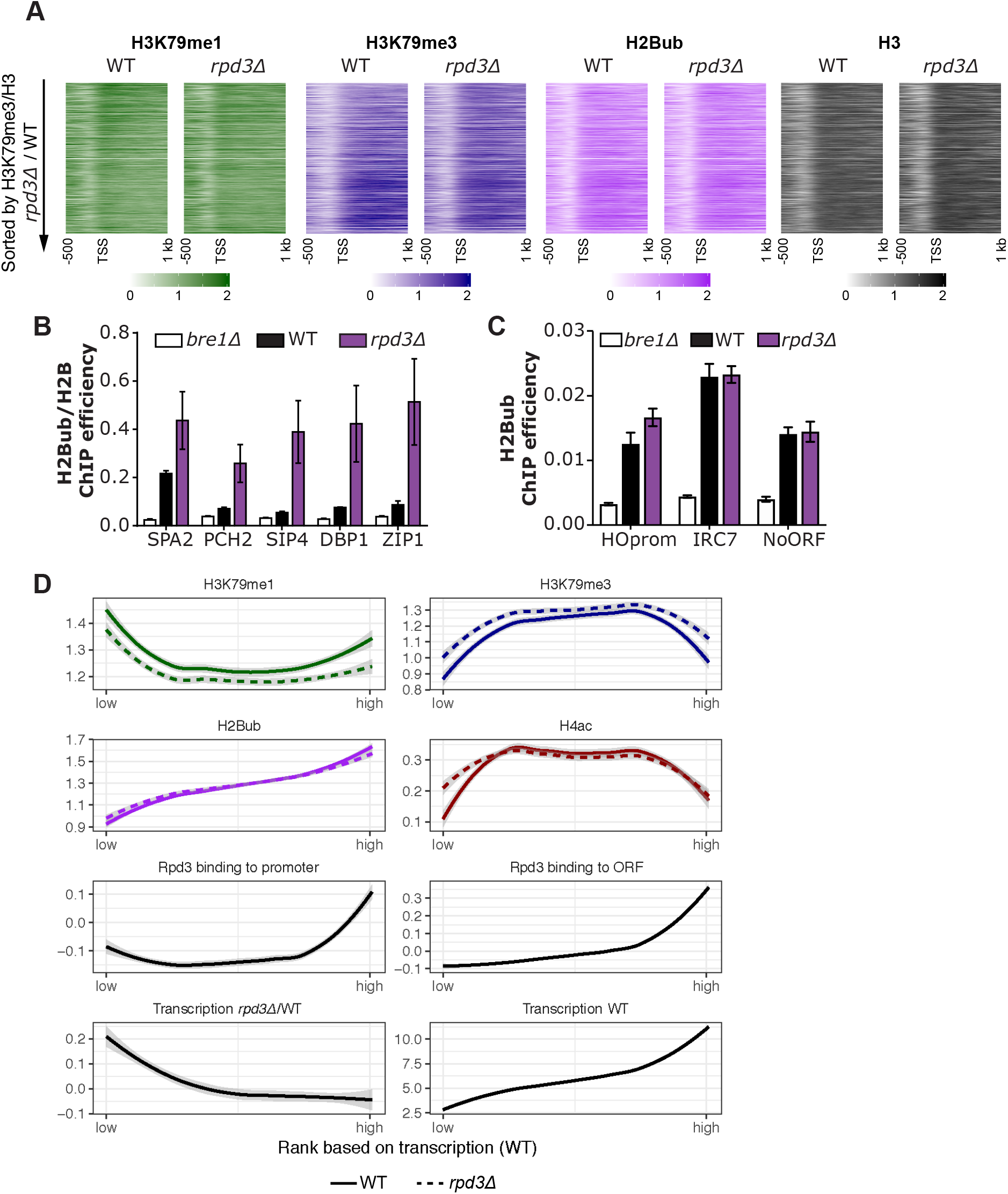
ChIP-seq and ChIP-qPCR data from WT vs *rpd3Δ* cells and the relation between transcription and histone modifications. A) Heatmaps of H3K79me1, H3K79me3, H2Bub and H3 sorted on H3K79me3/H3 rpd3Δ/WT. B) H2B-normalized H2Bub ChIP-qPCR IP efficiencies at several pre-selected loci in wild-type and *rpd3Δ* cells, with *bre1Δ* cells serving as a negative control. C) H2Bub ChIP-qPCR IP efficiencies at several pre-selected loci in wild-type and rpd3Δ cells, with *bre1Δ* cells serving as a negative control. D) ChIP-seq and RNA-seq data for genes ranked on transcription level in wild-type cells, smoothed using locally weighed regression. The shaded band around the line shows the 95% confidence interval. For H3K79me1, H3K79me3, H2Bub, H4ac and Rpd3 binding to ORF, the average coverage in the first 500 bp was used. Rpd3 binding to promoter was the average coverage in the 400 bp upstream of the TSS. Transcription in wild-type cells

**Supplemental Figure 3.**
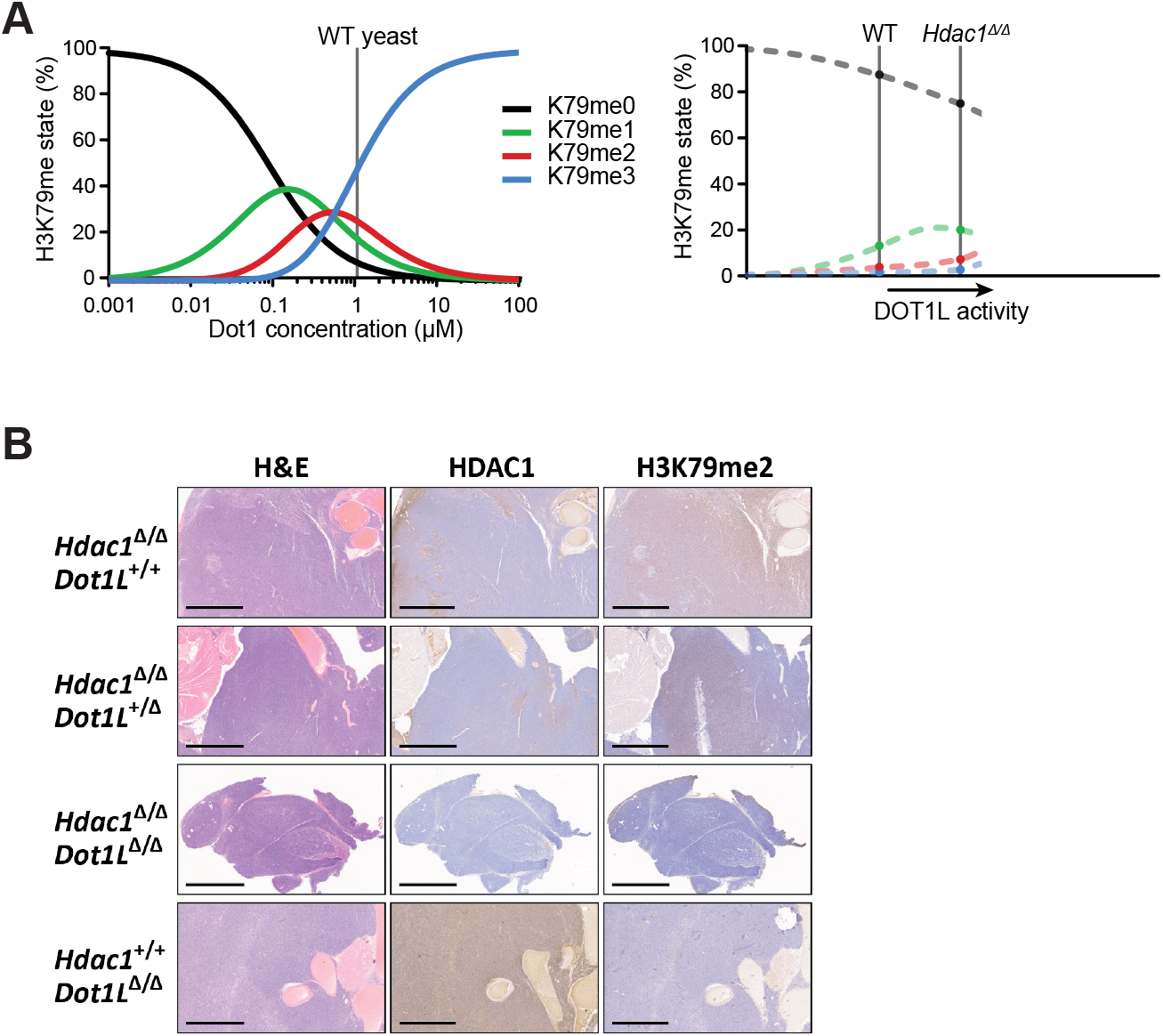
The distributive nature of Dot1, and thymus immunohistochemistry. A) Yeast Dot1 is a distributive enzyme and shows waves of the different methylation states over a range of Dot1 concentrations in yeast (De Vos et al. 2011), figure from De Vos et al. (2017). The catalytic nature of mammalian DOT1L enzymes is not known, but the observation that the abundance of each methylation state increases upon *Hdac1* deletion does not conflict with a distributive nature. Since H3K79 methylation levels are low in mammalian cells, it is possible we are looking at the start of the methylation waves, as indicated by the dotted lines. B) Representative H&E and immunohistochemical stainings of thymic lymphomas of the indicated genotypes. The scale bar represents 2 mm.

**Supplemental Figure 4.**
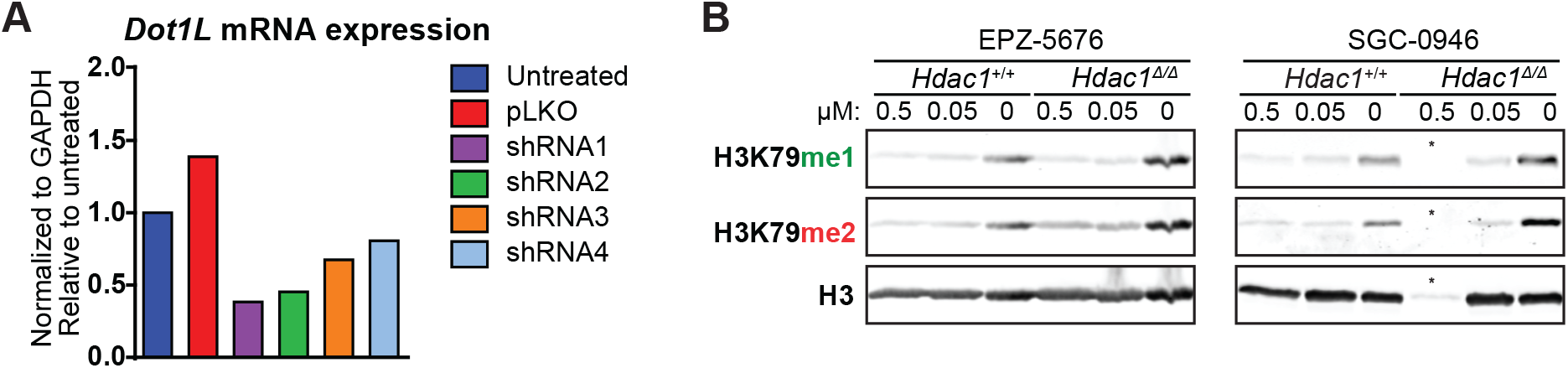
Control experiments belonging to Figure 4. A) Knockdown efficiency of the shRNAs used in Figure 4D. The shRNAs 1 and 2 give a knockdown of more than 50%. B) Immunoblot analysis of H3K79 methylation levels in the two cell lines with and without inhibitors. Both the effect of the inhibitors and the difference between the cell lines can be observed. * indicates a lane that was underloaded because most cells had died.

### Supplemental Methods

#### Mouse generation and crosses

The Dot1l^tm1a(KOMP)Wtsi^ mouse was generated by the Wellcome Trust Sanger Institute (WTSI) and obtained from the KOMP Repository (www.komp.org) (Skarnes et al. 2011). Since this mouse had a knock-out first allele, FLPe in a C57BL/6 background was crossed in to remove the FRT-flanked cassette (B6.Cg-Tg(ACTFLPe)9205Dym/A, MGI:2448985; Rodríguez et al. 2000). FLPe was crossed out to obtain C57BL/6 *Dot1L*^f/f^ mice, with LoxP sites flanking exon 2 of *Dot1L*. No thymic lymphomas were observed in C57BL/6 *Lck*-Cre;*Dot1L*^f/f^ mice. For this study, mice with the conditional *Dot1L* allele were crossed with mice bearing *Lck*-Cre and conditional Hdac1 alleles, which were described before (Heideman et al. 2013). All mice analyzed in this study were progeny of this cross and were in a mixed FVB/n, C57BL/6, and 129/Sv background. Wild-type control mice were the *Lck*-Cre-negative littermates of the other mice used in this study.

#### Nuclear extract preparation and immunoblotting

Thymic lymphoma cell lines were collected and washed by PBS. Single-cell suspensions of thymuses were obtained by passing the tissues through a 70 μm cell strainer, and cells were pelleted and washed with PBS. Samples were kept cold at all times and all buffers were supplemented with Complete protease inhibitors (Roche), Trichostatin A and nicotinamide. To make nuclear extracts, cells were first incubated in hypotonic lysis buffer (10mM Tris (pH 7.8), 5 mM MgCl_2_, 10 mM KCl, 0.1 mM EDTA, 300 mM sucrose, 5 mM B-glycerol) for ten minutes. Nonidet P-40 was added to an end concentration of 0.12% to rupture the cells. Nuclei were collected by centrifugation and lysed in RIPA buffer (20mM Tris (pH 7.5), 150mM NaCl, 1% Nonidet P-40, 0.5% sodium deoxycholate, 1mM EDTA, 0.1% SDS) for 30 minutes. All buffers were supplemented with Complete protease inhibitors (Roche), Trichostatin A and nicotinamide. Samples were sonicated for 2.5 minutes (10 second pulses) using the Diagenode Biorupter to solubilize chromatin. After this step, debris was pelleted and the supernatant was collected. Protein concentration was determined using the *DC* protein assay (Bio-Rad). The immunoblotting procedure was as described in (Vlaming et al. 2014). Yeast extracts were loaded on 16% polyacrylamide gels; murine extracts were loaded on gradient gels (4-12% Bis-Tris NuPAGE mini gels).

#### Antibodies

Blots with yeast samples were probed with antibodies against Dot1 (RRID: AB_2631109; Van Leeuwen et al. 2002), Pgk1 (A-6457, Invitrogen) and H2B (39238, Active Motif). Blots with mouse samples were probed with antibodies against HDAC1 (NB100-56340, Novus Biologicals), H3K79me1 and H3K79me2 (RRID:AB_2631105 and AB_2631106;Frederiks et al. 2008), H2BK120ub (#5546, Cell Signaling Technology), H3K9ac (ab4441, Abcam), total H3 (ab1791, Abcam), and a newly generated H4 antibody. Rabbit polyclonal antibodies against histone H4 were generated by immunizing with the peptide (C)VYALKRQGRTLYGFG of the C terminus of histone H4 of *S. cerevisiae*. The polyclonal serum recognizes human and mouse histone H4. ChIP experiments were performed using antibodies against H3K79me1, H3K79me3 and total H3 (RRID:AB_2631105, AB_2631107, AB_2631108; Frederiks et al. 2008) and antibodies against H2B (39238, Active Motif) and a new site-specific antibody against yeast H2Bub that we recently developed (Vlaming et al. 2016). For immunohistochemistry, antibodies against HDAC1 (ab31263; Abcam) and H3K79me2 (RRID:AB_2631106; Frederiks et al. 2008) were used.

#### ChIP-seq library preparation and data analysis

Library preparation and sequencing were performed by the NKI Genomics Core Facility, in two batches. Libraries from the ChIP samples from WT #1 were prepared using the TruSeq^®^ DNA LT Sample Preparation kit (Illumina, cat no. FC-121-2001), using ten times less adapter in the adapter ligation step. After fifteen PCR cycles, a size selection cleanup was performed using 0.5X Agencourt AMPure XP PCR Purification beads (Beckman Coulter, cat no A63881) to get rid of large fragments due to crosslinking DNA. The supernatant of the 0.5X cleanup was used to catch the smaller fragments; this supernatant was cleaned up 2 times with 1X beads to remove primers present in the libraries. Samples were pooled equimolarly and subjected to sequencing on an Illumina HiSeq2000 machine in a single-red 50bp run. Libraries from the ChIP samples from WT #2 and two *rpd3Δ* replicates were prepared using the KAPA HTP Library Preparation Kit, Illumina^®^ platforms (KAPA Biosystems KK8234), using Illumina-provided adapters at 200nM. After eleven PCR cycles, cleanup and pooling was as described above. Samples were sequenced in a single-read 65bp run on an Illumina HiSeq2500 machine. ChIP libraries from an *sin3Δ* strain prepared in the first batch (with WT #1) gave results comparable to the two *rpd3Δ* replicates from batch two. Reads were mapped to the *Saccharomyces cerevisiae* reference genome R64-2-1 with BWA version 0.6.1 and filtered for mapping quality 37 (Engel et al. 2014; Li and Durbin 2009). Each read was extended to 150 bp. Each sample was normalized for the sequencing depth by converting to Reads per Genomic Content (RPGC) with DeepTools (Ramírez et al. 2016). This was done by dividing the coverage by the sequencing depth, calculated as (total number of mapped reads * fragment length) / effective genome size (12.1×10^6^ bp). Data from the biological duplicates was found to be similar and the data sets were merged for further analyses.

#### Epi-ID analysis

Data from the Epi-ID H3K79me regulator screen described in Vlaming et al. (2016) were used. As described, a growth-corrected methylation score was calculated by first calculating the H3K79me3/H3K79me1 ratio and then subtracting the value expected based on the growth rate of the strain (Vlaming et al. 2016). The CLIK tool (Dittmar et al. 2013) was used to define groups of candidate regulators, and to determine the enrichment of the Rpd3L complex. Data on all components of Rpd3L and Rpd3S was obtained in the screen. Deletions were checked by PCR and barcodes were checked by Sanger sequencing. All deletions could be confirmed, with the exception of *sds3Δ*, which was eliminated from the plot in Figure 1B.

